# Gated electron transport in rhodopsin and its relevance to GPCR activation

**DOI:** 10.1101/650531

**Authors:** Angela S Gehrckens, Andrew P Horsfield, Efthimios M C Skoulakis, Luca Turin

## Abstract

We identify, by density-functional theory calculations, an electron donor-bridge-acceptor (DBA) complex within the highest resolution X-ray diffraction structures of rhodopsin. The donor is a conserved tryptophan, the acceptor is a zinc ion surrounded by a tryptophan, a histidine and a conserved glutamate. The unusual environment of the zinc ion confers high electron affinity on the zinc site. The bridge is the retinal which can exist either in the neutral aldimine (Schiff’s base) or aldiminium (protonated) state. When the retinal is unprotonated, no electron transfer occurs. Upon protonation of the aldimine, the DBA complex conducts and a full electron charge is transferred from donor tryptophan to the zinc complex. This gated electron transfer creates the molecular equivalent of a tunnel triode. Since rhodopsin is the ancestor of GPCRs, we discuss the possible relevance of this gated electron transport to other GPCRs, in particular to olfactory receptors which have been proposed to use an electron tunneling mechanism to detect molecular vibrations.

## Introduction

In 1996 Turin proposed that olfactory receptors detected molecular vibrations by inelastic electron tunnelling, i.e. electron transfer from a donor *via* an odorant to an acceptor, enabled by excitation of molecular vibrations in the odorant^1^. The physics behind this idea was shown to be sound^2-6^, and strong behavioural and perceptual evidence in its favour has come from experiments on insects and humans^7-10^. However, the idea currently remains controversial^11-13^, in part because vertebrate olfactory receptors are members of the G-protein-coupled receptor (GPCR) family: GPCRs are thought to work by a conventional lock and key mechanism^14,15^. Indeed, direct evidence of electron transfer in olfactory receptors is lacking. The most common objection to vibrational olfaction is therefore: “why should olfactory receptors be so different from other GPCRs?”^13^

GPCRs are derived from the evolutionarily ancient light-sensing protein rhodopsin and have evolved into ion pumps, ion channels and a variety of receptors sensitive to a vast range of ligands (see for example ^16^ for a review of the diversity of human GPCR receptors). It could therefore be argued that one more type of receptor activation mechanism would not be all that unusual. But this does not address a genuine physical problem which underlies objections to the vibrational theory of olfaction: proteins are typically wide-bandgap (4-5 eV) materials^17,18^. Charge carriers, be they electrons or holes, require energies much larger than thermal energy to be created. This energy is usually supplied by photons, whereas our noses work in the dark. So *how do electrons flow in olfactory receptors?*

We now address this question from a novel, more general perspective. Electron transfer requires a donor highest occupied molecular orbital (HOMO) to be higher in energy than an acceptor lowest unoccupied molecular orbital (LUMO): this describes a system temporarily out of equilibrium. Clearly, amino acids side chains alone cannot do this, since the side chain with the lowest LUMO energy (protonated lysine) is higher than the highest HOMO (tryptophan)^19^. We therefore postulated the existence of a metal ion in olfactory receptors, likely zinc bound to a histidine. The positive charge on the ion attracts electrons and thus lowers the LUMO energy to make electron transfer thermodynamically favorable. Zinc was proposed as a candidate for this mechanism in olfactory receptors, because of its known involvement in olfaction and the presence of several possible zinc-binding sites in olfactory receptors^1,20,21^. More recently copper has also been shown to be involved in odorant binding in olfactory receptors detecting thiols^22^

No GPCR olfactory receptor structures have been published to date. Remarkably, however, an intramembrane zinc ion has been found in x-ray structures of bovine rhodopsin, an evolutionarily ancient 7-TM protein thought to be the ancestral GPCR^23,24^. We recently proposed^5^ that some key components of the electron pathway required for olfaction are therefore already present in rhodopsin: a conserved tryptophan (donor) and metal complex (acceptor) on either side of the retinal. Here we study the energetics of electron transfer in rhodopsin in greater detail with density functional theory (DFT), using the highest-resolution published structure (pdb 1u19) to date.

## Zinc and electron transfer

Zn^2+^ has a high charge-density (≈112 C/mm^3^) ^25,26^. It coordinates readily to nitrogen, oxygen, and sulfur atoms. Its typical effect on electronic energy levels is illustrated in figure 1. While Zn^2+^ itself has no other available oxidation states, its charge can lower the energies of LUMOs in neighboring amino acids.

**Figure 1:**
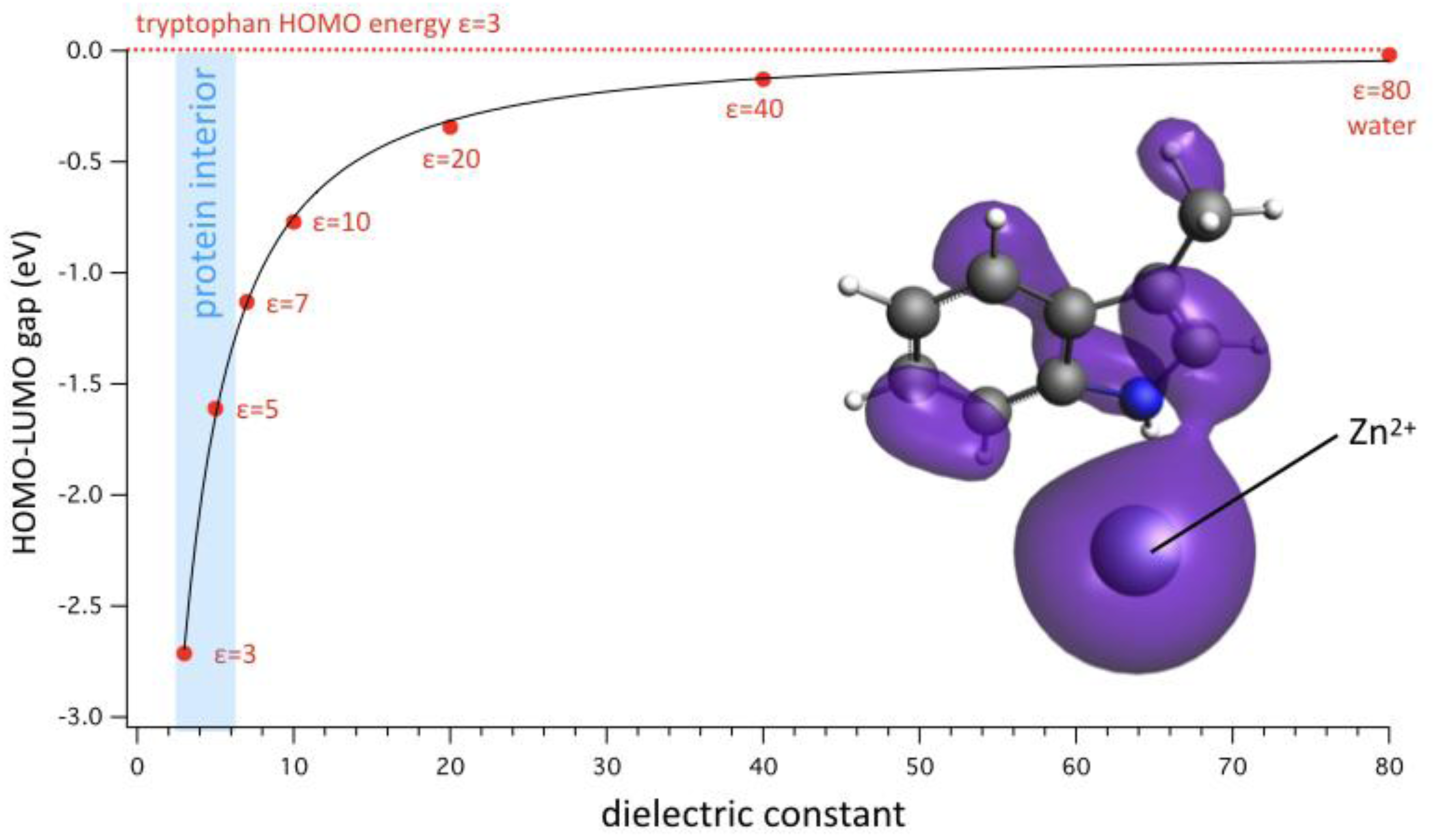
Computed energy gap between the HOMO of the tryptophan side-chain (methylindole) in a medium of ε=3 and the LUMO of methylindole coordinated with a Zn^2+^ atom in media of varying dielectric constant. Graph abscissa: dielectric constant. Ordinate: HOMO-LUMO gap. The gap is negative at all dielectric constants, showing that the presence of a coordinated zinc charge favors electron transfer from Trp to Zn^2+^-Trp. The exact energetics of the transfer will depend on the environment of the donor Trp, but transfer will be easier the higher the ε around donor Trp. The inset shows the Trp-Zn^2+^ complex and its LUMO in purple, whose shape and volume vary little as a function of ε. The spread of the LUMO over the fused rings of the tryptophan side chains suggests that part of the lowering of LUMO energy is due to charge delocalisation in the aromatic rings. The light blue region at left indicates the range of dielectric constants typically found in protein interiors. All calculations B3LYP-DZP with COSMO solvent, radius 2Å.

The data shown in figure 1 illustrate the energetics of this process. The presence of a coordinated zinc ion lowers the LUMO (in this case, of a Trp side chain) to below the energy of the donor Trp HOMO, and electron transfer is therefore energetically possible, provided that ionisation of the donor Trp does not incur too high an energy cost. The exact energetics will be determined by the environments of both donor and acceptor. Increasing the dielectric constant screens the charge and reduces the driving force. Coordination to an aromatic amino acid delocalizes the electrons and adds to the driving force.

The intramembrane zinc ion in rhodopsin was predicted from mutations^23^ and observed in the X.ray structure^27^ in 2004, i.e. after most of the major advances in rhodopsin structure and photochemistry were made, and has attracted relatively little attention since. Several features of this Zn^2+^ ion are unusual and initially controversial^24^: First, it is in the “wrong place”, approximately halfway across the membrane, and on the outer edge of the rhodopsin seven-transmembrane helix core. In other words, the environment of this charged ion will be mostly of low dielectric constant, which will do little to reduce its electrostatic self energy. Second, it is not present in all the published rhodopsin structures. Of the 5 structures of bovine rhodopsin we examined, for example, only 2 have the zinc ion present (figure 2). While there is no doubt that when present, Zn^2+^ is intrinsic to the protein, it seems to be labile during crystallisation, possibly due to the use of strong metal-chelating agents intended to inhibit metalloproteases.

**Figure 2:**
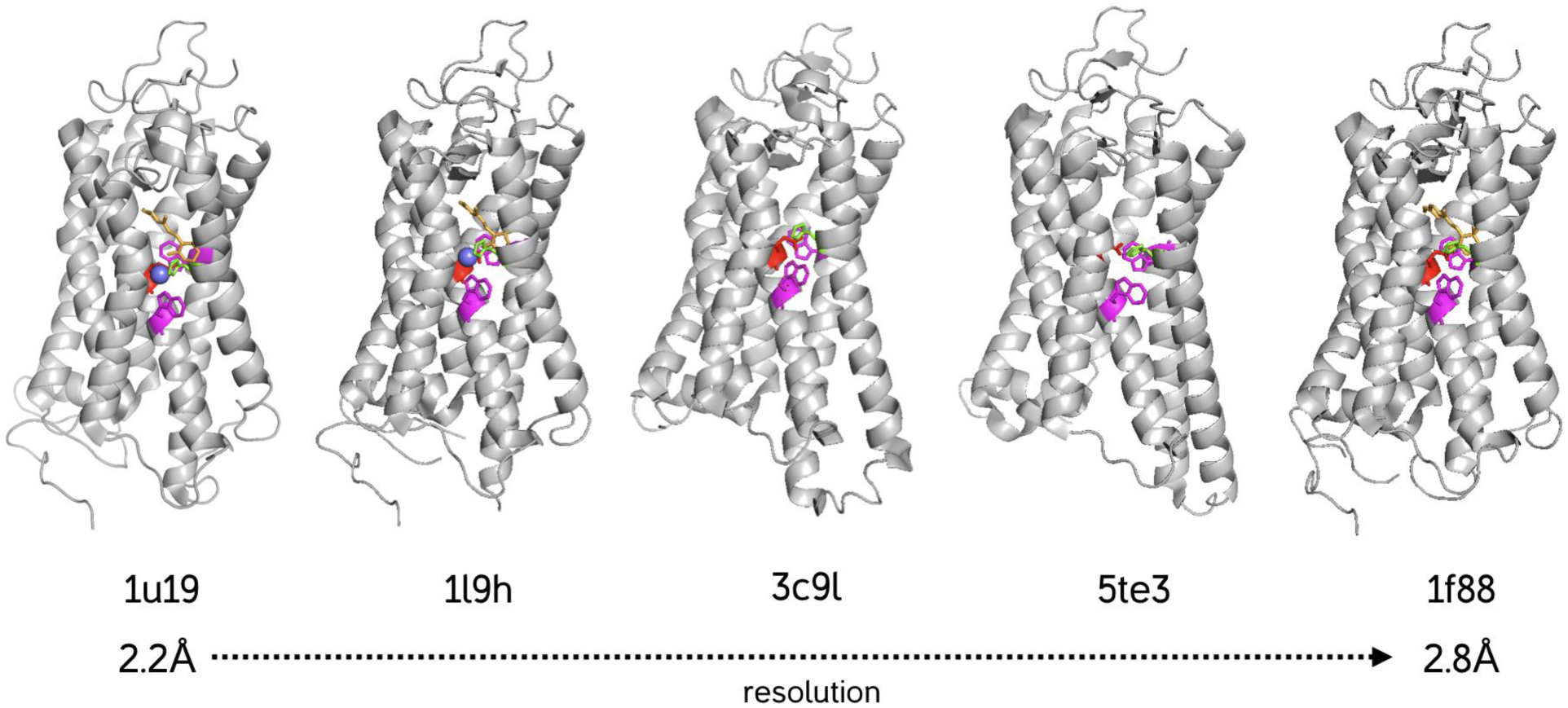
Structures of five bovine rhodopsins and opsins from the RCSB database 1u19 ^27^ 1l9h ^28^ 3c9l ^29^ 5te3 ^30^ and 1f88 ^31^. Only the two highest resolution structures contain the intrinsic Zn^+2^ ion inside the protein. The zinc ion is shown as a blue sphere, retinal in orange, tryptophan in magenta, glutamate in red and histidine in green.

## The environment of the Zn^2+^ ion

Zinc typically binds to one or more histidines and/or glutamates, and indeed those two amino acids are present in the vicinity of the rhodopsin zinc. The geometry of the site, however, is unusual. The typical geometry of zinc-histidine binding involves coordination of a deprotonated histidine nitrogen with the metal ion, and always results in the nitrogen pointing straight at the metal ion. In other words, the plane of the histidine ring should bisect the Zn^2+^. Similarly, carboxylic acids bound to zinc are typically deprotonated and the anion then interacts symmetrically with the metal ion, with the carboxylic carbon, the two oxygens and the metal forming a diamond shape^32^. Neither is the case in the two published high-resolution zinc binding site structures (figure3).

**Figure 3:**
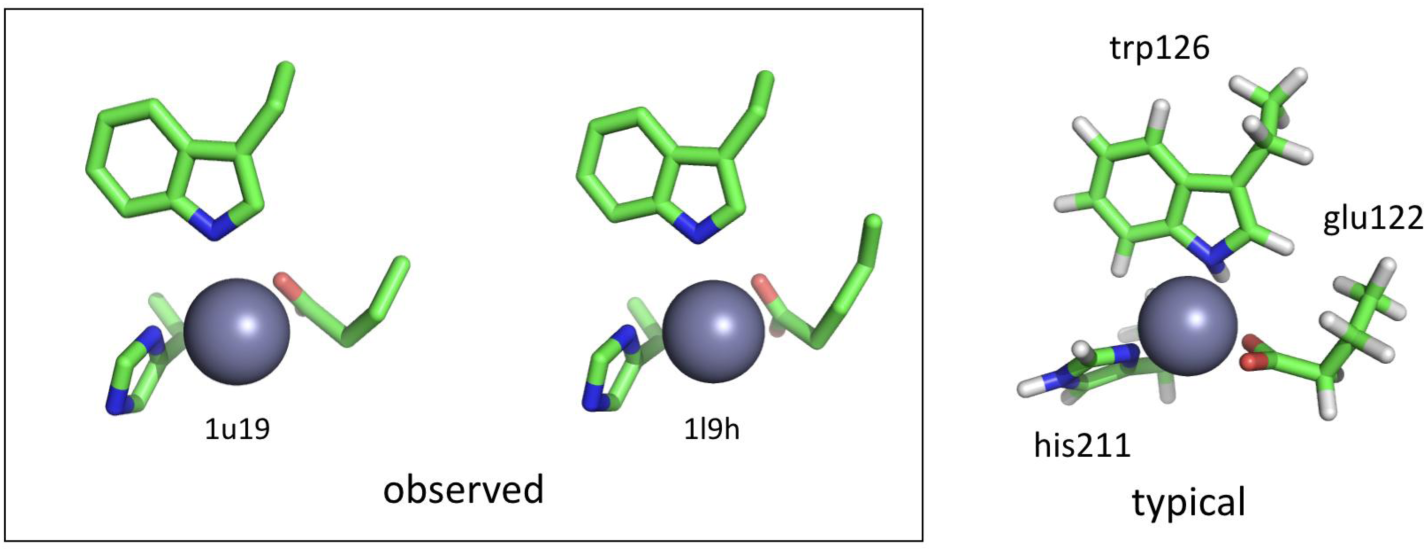
Structures of observed Zn-binding geometries in 1u19 and 1l9h (left) compared to the geometry obtained on the assumption of typical histidine-zinc binding via unprotonated ring nitrogen and symmetric coordination of the deprotonated carboxylate anion. Calculation: PBE-DZP geometry optimisation with alpha carbons fixed, ε=4.

**Figure 4:**
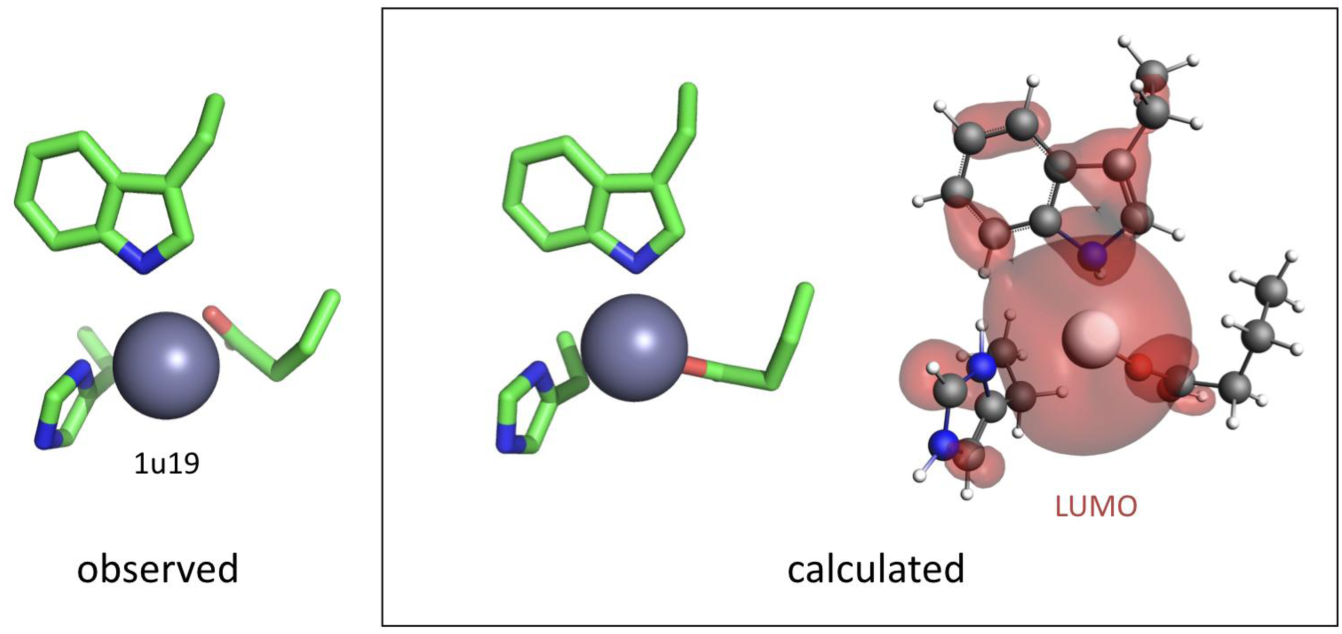
Structures of observed Zn-binding geometry in 1u19 (left) compared to the geometry calculated on the assumption of protonation of both histidine nitrogen and carboxylate (middle and right). The two calculated strutures have identical atom positions. Only heavy atoms are shown at middle for clarity, full structure and delocalised LUMO are shown at right to highlight electronic structures. Geometry optimisation B3LYP-DZP with alpha carbons fixed, ε=4.

The other two amino acids in the neighborhood of the Zn are tryptophan and a highly conserved glutamic acid residue. Tryptophan is a very weak acid and cannot bind to the Zn^2+^ via a deprotonated nitrogen. Indeed the geometry of the tryptophan with respect to the Zn^2+^ is similar to that of the histidine, tilted away from the Zn^2+^. The glutamic acid is known from spectroscopic results to be protonated^33^, i.e. uncharged, which is consistent with its geometry with one (carbonyl) oxygen pointing towards the Zn^2+^ and the other away from it. We have calculated the geometries of all possible combination of protonated and deprotonated amino acids. The configuration in which Trp and Glu are neutral, and His is cationic is the only one that matches the observed geometry (figure 5).

**Figure 5:**
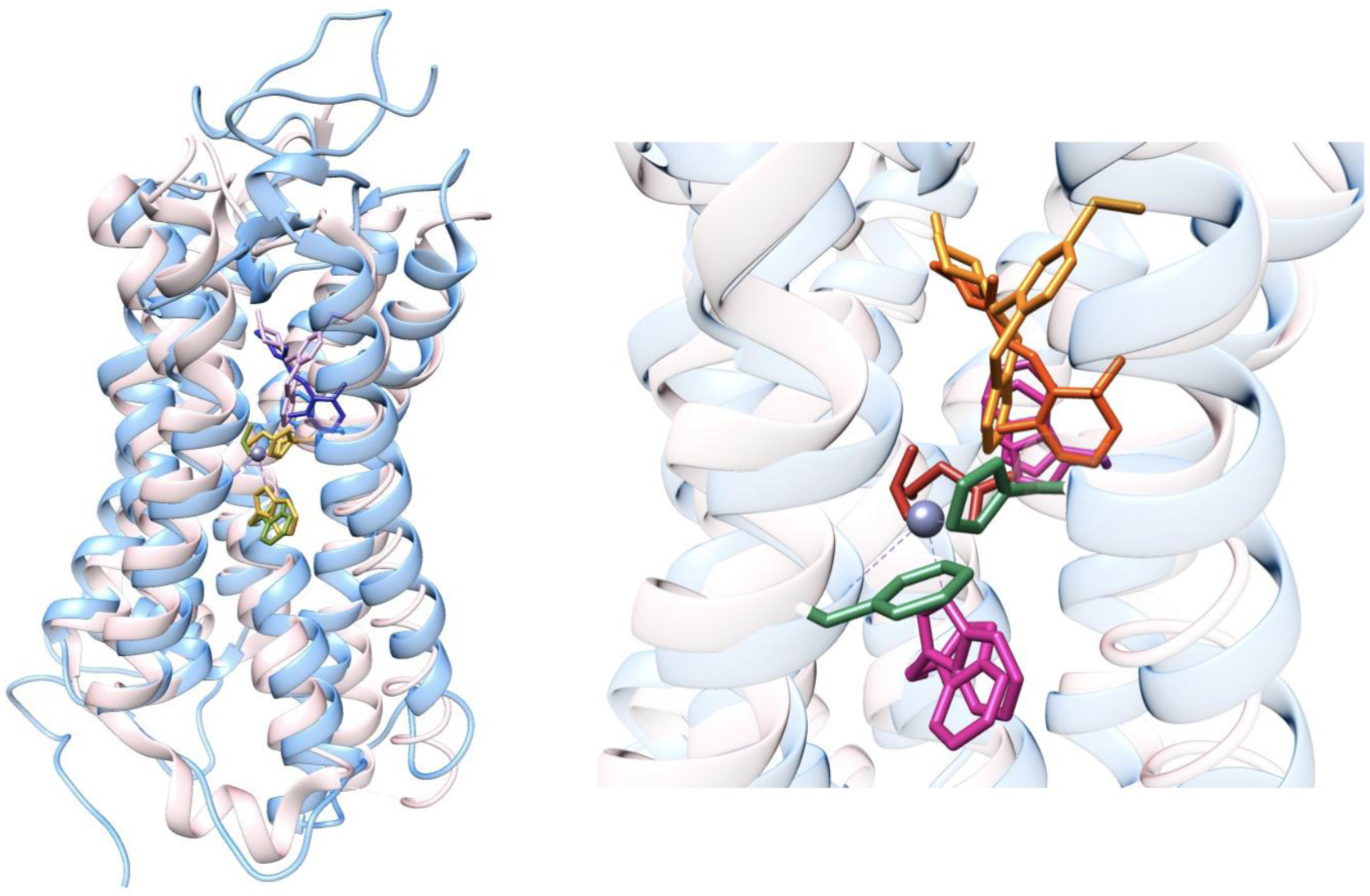
**Left:** Superimposed structures of rhodopsin (1u19)^27^ and human 5HT1B serotonin receptor (5v54)^36^ showing the remarkable similarity between helix positions, position of the acceptor tryptophan (Trp138 in 5v54) and the two ligands, retinal (blue) and metitepine (pink) (CAS Number 20229-30-5). The zinc in 1u19 is indicated. The structures were superimposed using the Mustang algorithm^37^ within the Yasara code. **Right:** close-up view of 1u19 and 5v54 showing donor and acceptor tryptophans (magenta), retinal and metitepine in two shades of orange, glutamates in red and His 211 of 1u19 together with Phe 176 of 5v54 in green.

One is therefore faced with three puzzling features of the zinc “binding” site: 1-No mitigation of the Zn^2+^ charge by counterions, 2-If anything, the electrostatic situation is worsened by the presence of the protonated histidine cation and 3-The lack of obvious coordination bonds between Zn^2+^ and its neighbors suggesting that the metal atom cannot play its customary structural role of bringing amino acids together^34^, and is in fact not tightly bound, which would explain its absence in most rhodopsin structures. The implications of this peculiar geometry of the zinc pocket (we suggest this is a preferable term to “binding site” since there seems to be little binding proper) will be discussed below in the context of electronic structure. For the moment let us note that the Zn^2+^ pocket contains an unusual concentration of unscreened, uncoordinated charge and aromatic groups, both features conducive to making a good electron acceptor site. The difference in LUMO energy between the calculated structure in figure 4 right and the typical structure in figure 3 right is due to the reduction in charge from +3 to +1, and the coordination of Zn^2+^by histidine. Together, these effects increase LUMO energy by ≈ 5eV.

## The electron donor

The small, hydrophobic ligands (odorants) that activate olfactory receptors are in a sense intermediate between retinal, a large hydrophobic ligand, and, say, serotonin, an odorant-sized but hydrophilic ligand. Comparison of the structures of rhodopsin and a serotonin receptor with an inverse agonist bound shows remarkably little difference between the positions of these very different ligands (figure 5). With olfactory receptor function in mind, and given that the inelastic electron tunnelling mechanism will only work if donor and odorant are very close to each other, we are looking for a good electron donor amino acid on the other side of the ligand from the zinc region. An obvious candidate for this electron donor is the completely conserved tryptophan found in all GPCRs, numbered 265 (Trp 265^6.48^ in Ballesteros-Weinstein notation^35^) in rhodopsin. In the serotonin receptor it is in contact with the ligand (here metitepine), in rhodopsin it is in direct contact with retinal.

## The retinal bridge

Compared to GPCR ligands such as neurotransmitters, retinal exhibits several well-known peculiarities^38-41^. The first is a series of alternating conjugated double bonds. Second, retinal binds to opsin *via* an aldimine (Schiff’s base) linkage obtained by condensation of the retinal aldehyde with the amino group of a neighboring lysine. The aldimine is known to be protonated and stabilised by a nearby carboxylate, Glu 113. This has two effects: first, the HOMO-LUMO gap is greatly reduced by protonation, which redshifts the retinal absorption to around 580 nm. Second, a protonated aldimine is a strong electron acceptor, whose electron-withdrawing effect propagates via the alternating double bonds from one end of the molecule to the other. Retinal can therefore be considered a molecular wire whose electrical potential is controlled by aldimine protonation (figure 6). In what follows, we will be concerned with the effect of protonation and deprotonation on electron tunnelling at right angles to the retinal. In other words, we treat the retinal as a bridge between donor and acceptor, a bridge whose charge and electronic structure is altered by changing the protonation at the aldimine end.

**Figure 6:**
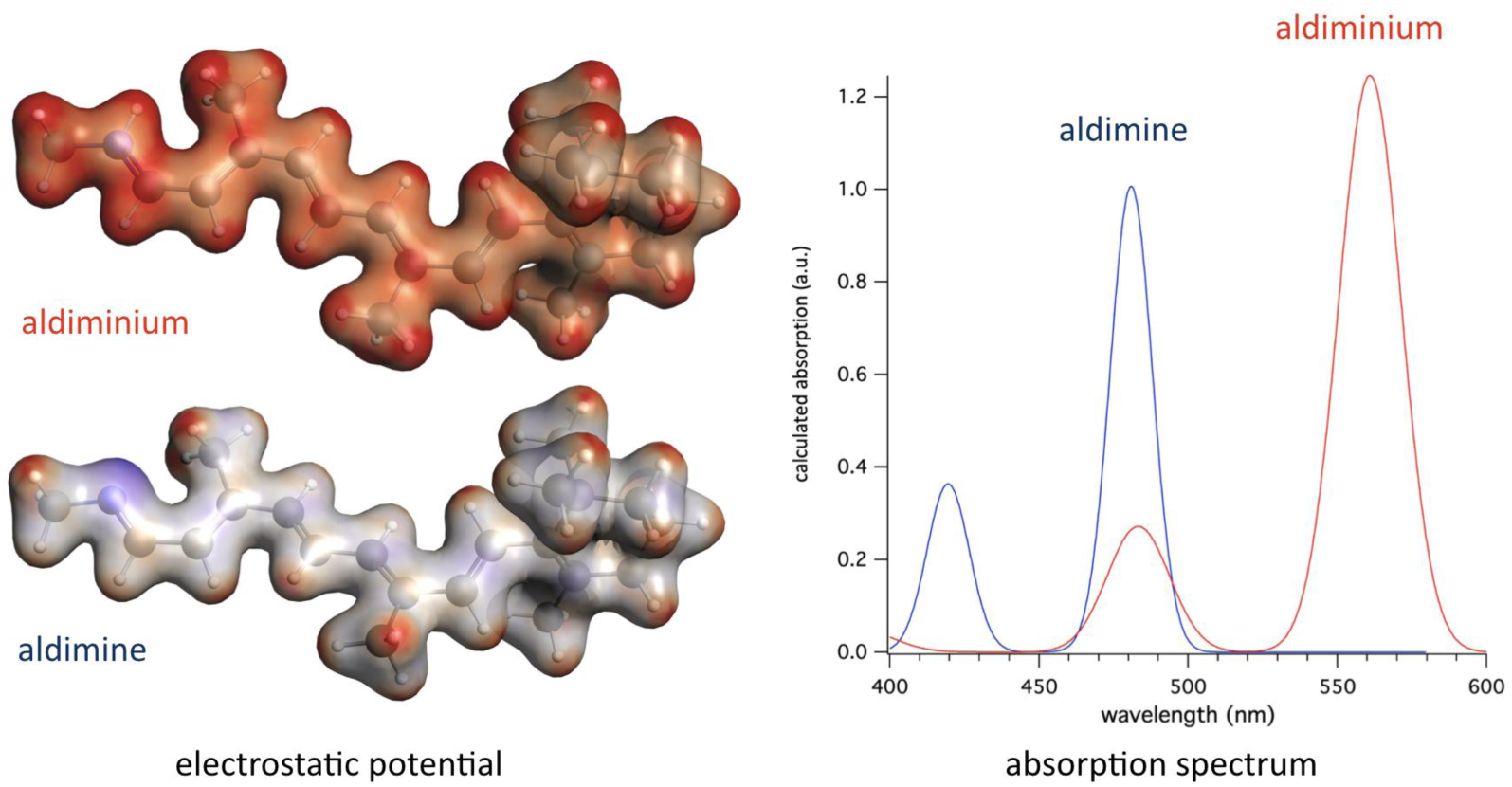
**Left:** Electron-density surface of all-trans retinal with the terminal aldimine in the protonated (top) and unprotonated (bottom) states. Electrostatic potential is mapped onto the electron density surface with the same boundary values in both cases. Red is positive. The presence of alternating double bonds allows the molecule to act as a molecular wire and the electron withdrawal by the aldiminium results in the entire molecule becoming positively charged. **Right:** Absorption spectra of aldimine (blue) and aldiminium calculated by Time Dependent DFT. The red-shift of the main absorption peak caused by protonation is clearly visible, to approximately 570 nm, close to the experimental value.

## Modeling of the Donor-Bridge-Acceptor complex

A DFT calculation of electronic structure in reasonable time requires paring down of the donor-bridge-acceptor complex to its essentials. We have retained the side chains including alpha carbons of four amino acids: the donor tryptophan and the three amino acids, namely His, Trp and Glu surrounding the zinc. The retinal is present either as an unprotonated aldimine or as a protonated one accompanied by its counterion Glu. All initial coordinates are taken from the 1u19 pdb file. We fixed the positions of the zinc, all alpha carbons and the two extreme carbons of the retinal (see figure 7). Starting from the pdb coordinates, the structure is minimised using a dispersion-corrected PBE-D3(BJ) functional and a double-zeta polarisation basis set (DZP). The charge is set to +3 (see discussion of the Zn^2+^ pocket structure above, i.e. two charges on the Zn and one on the histidine). Note that the bridge (retinal aldimine or aldiminium + glu counterion) is neutral in both cases since in one case the aldimine is neutral and in the other the aldiminium cation is balanced by the glu anion. The medium is assumed to have a dielectric constant equal to 4 ^42,43^ and a solvent radius of 2Å.

**Figure 7:**
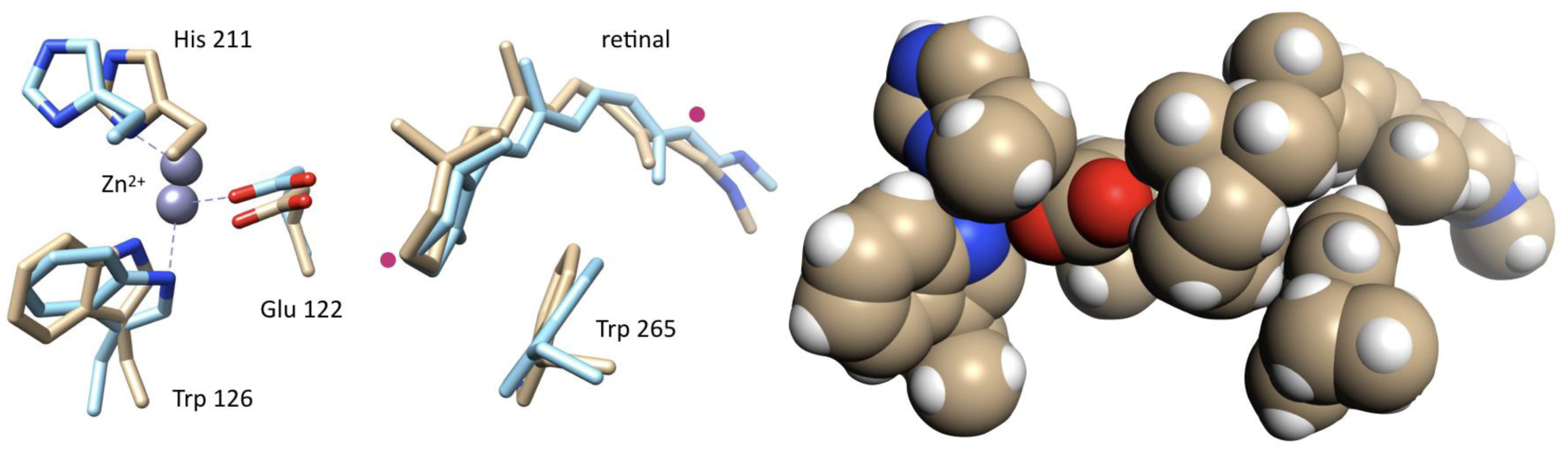
**Left:** Superimposed structures of the donor-bridge-acceptor complex from 1u19 (coordinates from pdb) in brown and the energy-minimised structure calculated by DFT by minimising energy with alpha carbons fixed on the four amino acids and two carbon atoms of the retinal (red dots). Unsurprisingly given the large change in charge and electronic structure, the largest changes are seen at the acceptor end (Zn^2+^ complex). The energy minimisation was done using B3LYP-DZP, a charge of 3 and ε=4 (COSMO model). **Right:** space-filling model of native 1u19 structure showing the close packing of the donor-bridge-acceptor complex.

Energy minimizations are done in four different configurations: 1-closed shell ground state for aldimine; 2-diradical state for aldimine, assuming complete transfer of one electron from donor tryptophan to zinc complex; 3-closed shell ground state for aldiminium; and 4-diradical state for aldiminium. The biggest structural changes between native and minimised structure occur for (4), and are shown in figure 3 right. They are surprisingly modest considering the different —classical—force fields used to fit the crystal structures to the electron densities. Comparison of the native structure extracted from 1u19 and the aldiminium diradical after minimisation is shown in figure 7.

## Energetics of electron transfer

The computed energetics of electron transfer are shown in figures 8 and 9 for the unprotonated and protonated aldimine respectively. When the aldimine is uncharged, both the system frontier orbitals are confined to the donor Trp and retinal in the ground state. The HOMO-LUMO gap is 1.7 eV, and the diradical state is *higher* in energy by 0.5 eV. When the aldimine is protonated to aldiminium, with the Glu counterion now present, the closed-shell HOMO-LUMO gap falls to 0.2 eV and the diradical state is now the ground state, energetically lower by 1.6 eV. The spin surface (sum of both unpaired electrons on donor and acceptor) now spans the entire DBA complex. In other words, protonation of the retinal aldimine has had a profound effect on electronic structure, in essence allowing the DBA complex to conduct electrons. This is tantamount to voltage control of conduction in a classic triode, except at the nanoscale, and resembles the tunnel triode arrangement proposed by Chang and Esaki ^44^ but never built because of physical constraints. In their words, “The difficulty is an obvious one: The region through which the carriers tunnel is “nonconductive”, and to attach a third control electrode to it is meaningless”. In our case, the retinal molecular wire conducts and acts as both base and control electrode, with a proton setti ng base voltage. Rhodopsin therefore also resembles a single-molecule ChemFET ^45^.

**Figure 8:**
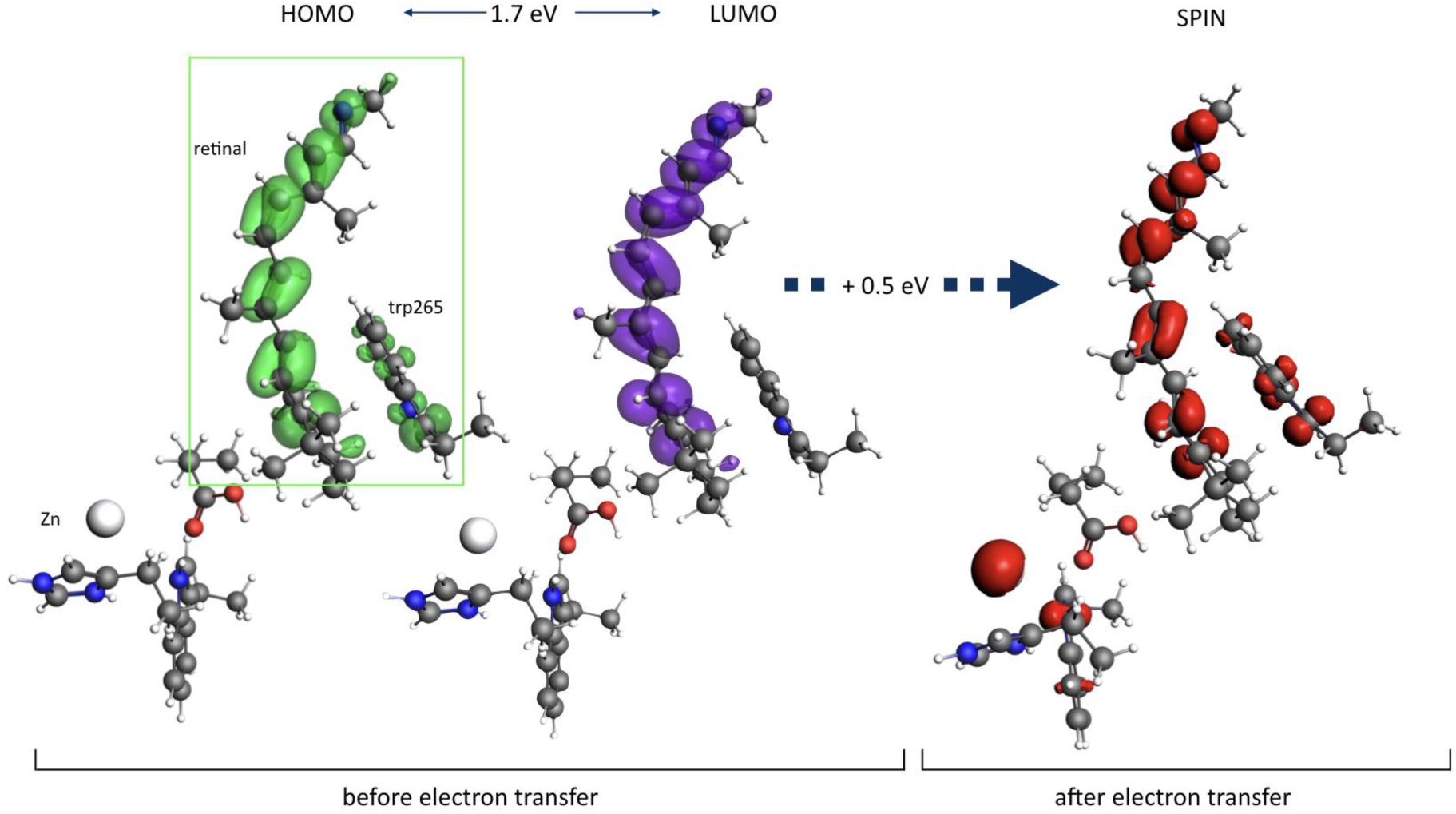
Electronic structure of the donor-bridge acceptor complex with unprotonated retinal aldimine. **Left**: complex in the ground state, with HOMO shown in green and LUMO in purple. Both HOMO and LUMO are confined to donor trp265 and retinal (inside the green rectangle). HOMO-LUMO gap is 1.7 eV. **Right:** the same structure after energy-minimization in the diradical state. Spin is now distributed between donor and acceptor sites, but the diradical state lies 0.5 eV (≈ 11 kcal/mole) above the closed-shell ground state. Calculations B3LYP-DZP with ε=4, COSMO model.

**Figure 9:**
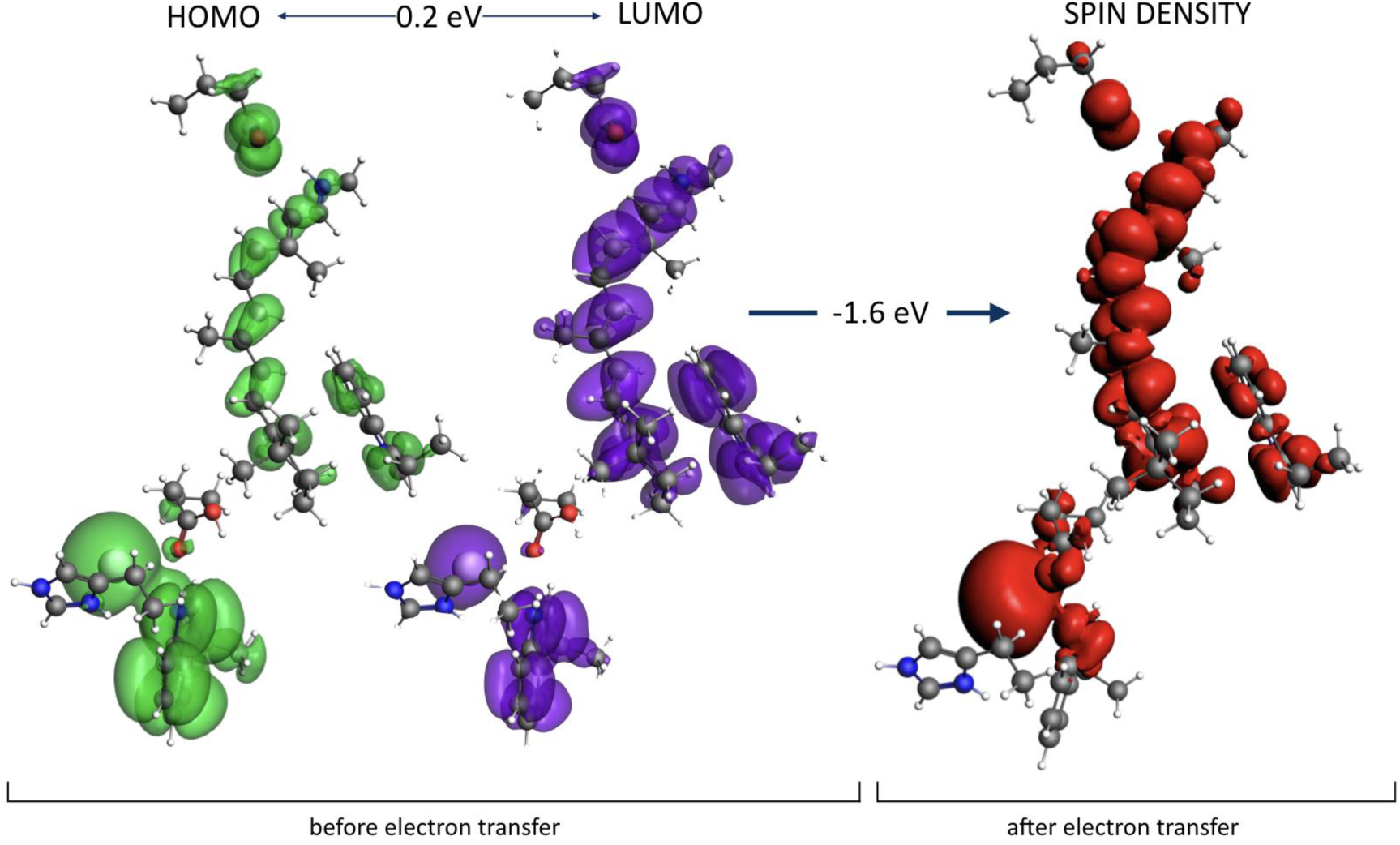
Electronic structure of the donor-bridge acceptor complex with protonated retinal aldimine. **Left**: complex in the ground state, with HOMO shown in green and LUMO in purple Both HOMO and LUMO are now spread throughout the DBA complex. HOMO-LUMO gap is 0.2 eV. **Right:** the same structure after energy-minimization in the diradical state. Spin is now evenly distributed between donor and acceptor sites, and the diradical state lies 1.6 eV (≈ 11 kcal/mole) below the closed-shell state, and is therefore the new ground state.

## Implications of an electronic receptor mechanism

The purpose of this study was to assess the plausibility of our proposal that vertebrate olfactory receptors are in fact electronic devices that use inelastic electron tunneling across the odorant to probe its vibrations. We wanted to see whether there were any structural indications of such electronic circuitry within GPCRs whose structure has already been determined. In this respect our search has been successful. We find in rhodopsin, the ancestor of GPCRs, an arrangement of electron donor and acceptor on either side of the retinal. If the structure in vivo is identical to its counterpart in crystals, it will allow electron transfer from a highly conserved donor tryptophan to a zinc pocket situated on the other side of the retinal ligand.

Zinc is unusual among transition metals found in biology in having no redox chemistry, and only existing in the +2 state. After electron transfer, therefore, the formal oxidation state of the Zn^2+^ is unchanged and the electron resides primarily on the neighboring tryptophan. As is shown in figure 4, among the different protonation states of the zinc pocket amino acids, only the one consistent with the crystallographic structure has an electron affinity sufficient to cause electron transfer. What happens to the electron after transfer? Our calculation suggests a high pH-sensitivity of the electron affinity. Deprotonation of the histidine, the glutamic acid or both will have two effects. First, it will turn the zinc pocket into a proper high-affinity zinc binding site with proper coordination to customary ligands of Zn^2+^ (figure 3 right). Second, provided the missing electron on the donor Trp has been replenished, deprotonation of His 211 and Glu 122 will raise the energy of the Zn^2+^ complex LUMO by approximately 5 eV (see figs 3 and 4), to a point where the electron will seek to travel onwards, possibly to effect a redox reaction ^1^.

The idea that the zinc pocket can exist in protonated and deprotonated states also offers an answer to the question of how the Zn^2+^ observed in rhodopsin is inserted in the protein in the first place. In the state in which the zinc pocket is observed in the crystal, inserting Zn^2+^ would be an energetically unfavorable process. Deprotonating the histidine and glutamic acid in the pocket, however, would allow Zn^2+^ to bind. If this deprotonation were temporary, it would then permit the bound Zn^2+^ to perform its electron acceptor function.

The effect of electron transfer in the rhodopsin model is to create a hole on the highly conserved donor tryptophan. It is reasonable to assume that, much as happens in other proteins^46,47^, this hole will be filled by hopping from other aromatic amino acids^48^. If the electron is ejected from the acceptor, possibly by protonation as described above, the electron transfer process can be repeated at will. Note that the rate of transfer need not be very high. Receptors work on millisecond time scales, and a flux of electrons of the order of 1-10/ms, i.e. of the order of a femtoampere, would suffice. Observed protein electron conductivities, for example as seen in the scanning tunneling microscopy^49^ and in photochemical studies^50^, can be orders of magnitude higher. The exact topology of the electron circuit, the nature of the ultimate electron donor and acceptor, respectively before and after the electron transit through the receptor, remain to be determined.

Our proposed electron transfer mechanism seemingly plays no first-order role in light transduction by rhodopsin. Activation of rhodopsin proceeds by isomerisation of the retinal followed by a complex sequence of steps that leads to activation of the receptor^40^. None of these involve electron transfer in an obvious fashion. What then could be the function of this arrangement of amino acids and metal? One possibility is that the arrangement of donor and acceptor orbitals is intended to influence the electronic structure of retinal and tune its absorption spectrum. Another is that the isomerisation of retinal interacts with electron transfer, possibly by modifying the energetics of aldimine deprotonation, known to be necessary to signal transduction^51^ and that this in turn has an effect on further steps in signal transduction. Both possibilities need to be investigated by more refined DFT calculations, possibly involving the whole protein^52^. It is not clear whether the zinc pocket in the rhodopsin structure seen in crystals is in its physiological state. If it is, charge transfer will create unpaired electrons and therefore electron spins, chiefly residing on the donor and acceptor tryptophans. This should give rise to an ESR signal, though it may be weakened by its rather broad distribution as seen in figure 9 (P J Hore, personal communication).

## Relevance to other GPCRs

We emphasize that we are not suggesting that this circuit is functional in rhodopsin, or that it plays an important role in its primary function, though mutations in amino acids in the zinc binding site, notably His 211, are associated with Class II retinitis pigmentosa^23,53^. We are instead proposing that the elements of electronic circuitry we observe in rhodopsin, where they serve no obvious function, could have evolved in other contexts to become essential. The specific electronic mechanism posited for olfactory receptors, which is to probe odorant vibrations by inelastic tunnelling, may be unique to olfaction and has been implemented in laboratory devices^54^. Olfactory receptors have unique requirements: they are intended to bind molecules never encountered before and to probe their chemical composition. Both requirements are fulfilled by relatively nonspecific receptors^55^ that perform vibrational spectroscopy. It is unlikely that such a mechanism would be of much use in other receptor systems, particularly in those, such as neurotransmitter receptors, where binding is highly specific.

In this context, the structure depicted in figure 5, 5v54^36^, of the human 5HT1B receptor bound to the antagonist metitepine, is of particular interest. Metitepine (methiotepine, PuBChem 4106) is a promiscuous antagonist acting on 5-HT, dopamine and adrenergic receptors. Its structure is tricyclic, distantly related to phenothiazines but with a central thiepine 7-membered ring. When bound to the receptor, metitepine has the conformation shown at left in figure 10. We were surprised to find that this conformation did not match the closed-shell geometry of the molecule calculated by DFT: the angle between the two benzene rings flanking the thiepine is less flat in the calculated conformation.

**Figure 10:**
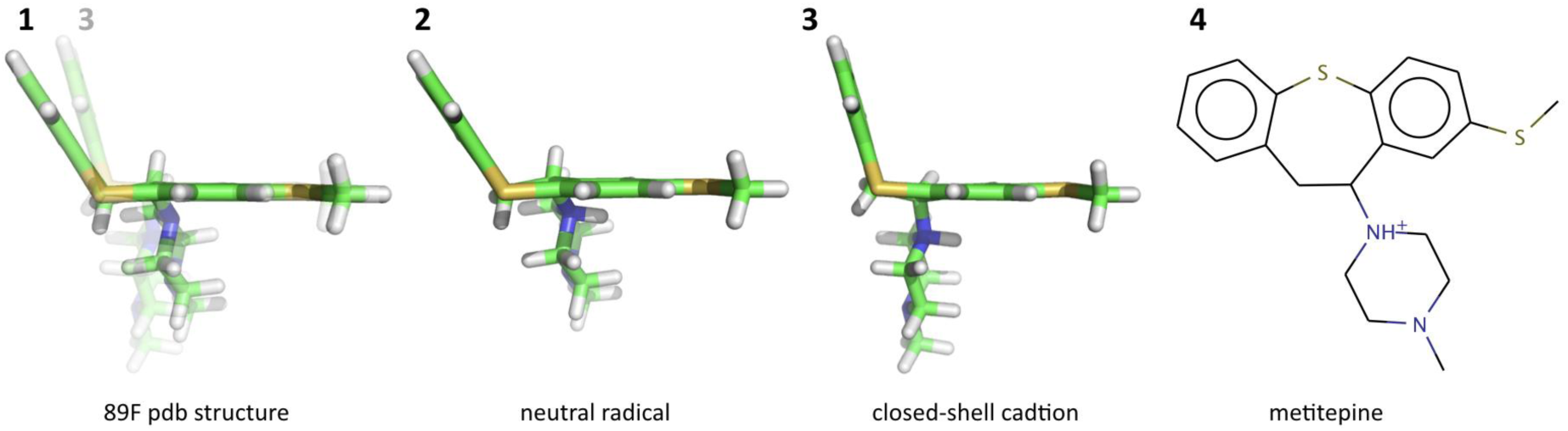
From left: **1:** structure of the metitepine (pdb 89F) found in the 5HT1B receptor (pdb 5v54). **2:** structure of the neutral radical calculated from the native structure by protonation of the piperazine ring nitrogen adjacent to the thiepine and addition of one electron. The neutral radical conformation closely approximates that of the pdb structure both in ring dihedral angle and position of the piperazine side chain. **3:** energy-minimised structure of the closed shell cation. This structure is also shown in transparency behind **1** to illustrate the structural differences. Forcing **3** into conformation **2** without adding an electron requires 0.96 eV (energies B3LYP-DZP in vacuo), or ≈38 times thermal energy at 300K. All geometry calculations use PBE-DZP. **4:** Structure of metitepine (1-methyl-4-(8-methylsulfanyl -5,6-dihydrobenzo [b][1]benzothiepin-6-yl) piperazine).

It seemed interesting to ask whether the conformation observed in the receptor might be that of a radical. A radical could form following electron gain by the metitepine cation, since the piperazine ring will be protonated at physiological pH. The structure of the neutral metithepine radical (electron gain from cation) gives a better match to the pdb structure, suggesting that receptor-bound metitepine contains an unpaired electron (figure 10). The resolution of 5v54 (3.9Å) is probably not sufficient to determine conclusively which structure is present, and the neutral radical shows some instability in the thiepine-nitrogen bond which would probably require stabilisation to avoid fission. Nevertheless, this surprising finding is in agreement with our proposal that GPCRs are electronic devices. A strong electron acceptor or donor bound to the active site could perturb electron transport either by acquiring an electron without passing it on to zinc, or facilitating electron transport by hole creation (electron loss). Either way, a receptor-bound drug in the radical form should be paramagnetic, and this property could be measured on receptor preparations bound to metitepine, provided the number of receptors exceeds the threshold of EPR detection, of the order of 10^9^ spins.

The 5v54 structure contains an acceptor tryptophan in the same position as that binding zinc in rhodopsin, though no zinc is present in the structure. We predict that if care is taken not to chelate it during purification and crystallization, this “missing zinc” will be found in this receptor, and likely in others. At the time of writing, 40 GPCR structures have been determined. The fact that metitepine binds to, and inhibits, a variety of receptors could be used to check whether this mechanism is common to different receptor types. Indeed, the existence of a large class of nonspecific activators known as pan-assay interference compounds (PAINS for short)^56,57^ many of which are electron donors and acceptors, may be more a gain than a pain for drug discovery. The well-known case of rhodanine^58^ (2-Sulfanylidene-1,3-thiazolidin-4-one) may be accounted for by the fact that it is a small, powerful electron donor. Its action may be due to interference with receptor activation, rather than to catch-all “redox cycling” and “free radical” mechanisms. If GPCRs turn out more generally to be electronic devices, then so will their ligands. The implication for GPCR pharmacology is that the electronic structure and affinity of the ligand may be of importance. Similar general ideas have been proposed before^59,60^ and the remarkable biological effects of electroactive molecules have been noted^61^. Electroactivity is correlated with function and used in the analysis of many useful drugs^62^.

In summary, in addition to the inelastic tunneling mechanism proposed for olfaction^5^, there may be other ways for a GPCR ligand to control electron flow. In the case of elastic tunneling, control over the height of the tunnelling barrier could be effected merely by the ligand bearing a positive charge to lower an empty orbital on the bridging molecule, which is of course the case for all biogenic amines and the vast majority of alkaloids (the term itself denotes a protonatable base). In the third case, a redox mechanism is used in which activation involves electron donation by the ligand itself. It may apply to PAINS, tricyclic drugs known to be electron donors like phenothiazines ^3,64^, related tricyclics like metitepine and possibly to catecholamines. Catecholamines are good electron donors (actually a hydrogen atom) and readily form semiquinones and quinones^65^. The electron affinity of catecholamines is controlled by the cationic group. Neutralisation of the amine by base in dopamine, for example, immediately leads to oxidation, formation of quinones and polymerisation into melanin^66^. The different mechanisms are shown in figure 11, redrawn from^67^. The three types of receptors illustrated are closely related to one another and rely in subtly different ways on a ligand-gated electron current. In conclusion, all the above suggests that olfactory (inelastic) receptors, far from being outliers as has been argued, may arise from minor evolutionary adaptations of the GPCR structure that enable them to detect odorant molecular vibrations.

**Figure 11:**
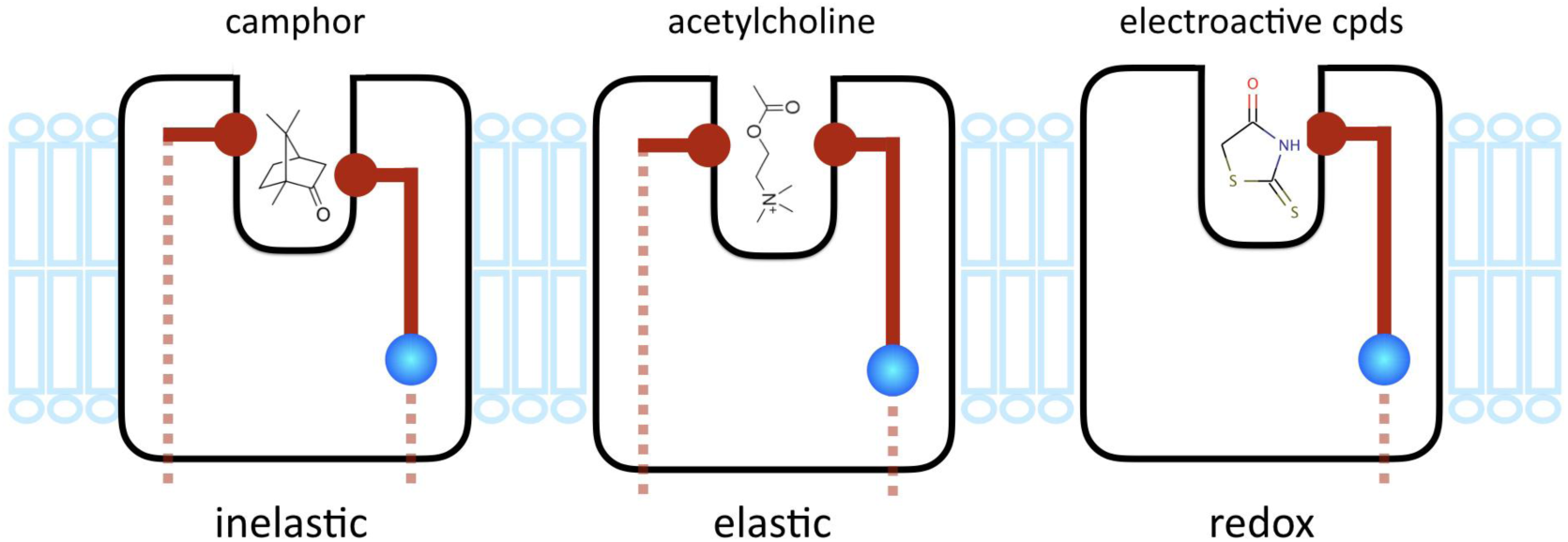
Proposed types of receptors using an electronic circuit to “read” a bound ligand. **Inelastic** receptors have a tunneling circuit traversing the receptor with an energy jump across the tunneling gap. To turn on such a receptor, the molecule has to be bound and possess one or more vibrations at the correct energy. **Elastic** receptors have the same circuit topology, but without an energy jump. The receptor is turned on when a molecule binds to it and includes a feature, such as a positive charge, that lowers the barrier to electron tunneling by lowering the energy of an empty bridge orbital close in energy to donor and acceptor values and acting as a channel for electron flow. **Redox** receptors have only the output half of the electron circuit. The electrons come from the bound molecule itself (rhodanine is depicted here), which undergoes an oxidation step when bound.

## Methods

DFT calculations used Amsterdam Density Functional code^68^ (ADF, www.scm.com) version 20199.102 and earlier, running either on a 12-core Mac Pro or on 50-120 cores at www.crunchyard.com. Protein model alignments were made using Yasara^69^ www.yasara.com.Structures were visualised with PyMol^70^ and Chimera^71^. Data were plotted using IgorPro (www.wavemetrics.com).

## Acknowledgments

LT thanks the Stavros Niarchos Foundation and DARPA for support. We thank the staff at SCM.com and Crunchyard.com for help with the DFT calculations. AG acknowledges the support of an Erasmus+ Scholarship.

